# Circumstantial Evidence? Comparison of Statistical Learning Methods using Functional Annotations for Prioritizing Risk Variants

**DOI:** 10.1101/011445

**Authors:** Sarah A. Gagliano, Reena Ravji, Michael R. Barnes, Michael E. Weale, Jo Knight

## Abstract

Although technology has triumphed in facilitating routine genome re-sequencing, new challenges have been created for the data analyst. Genome scale surveys of human disease variation generate volumes of data that far exceed capabilities for laboratory characterization, and importantly also create a substantial burden of type I error. By incorporating a variety of functional annotations as predictors, such as regulatory and protein coding elements, statistical learning has been widely investigated as a mechanism for the prioritization of genetic variants that are more likely to be associated with complex disease. These methods offer a hope of identification of sufficiently large numbers of truly associated variants, to make cost-effective the large-scale functional characterization necessary to progress genome scale experiments. We compared the results from three published prioritization procedures which use different statistical learning algorithms and different predictors with regard to the quantity, type and coding of the functional annotations. In this paper we also explore different combinations of algorithm and annotation set. We train the models in 60% of the data and reserve the remainder for testing the accuracy. As an application, we tested which methodology performed the best for prioritizing sub-genome-wide-significant variants (5×10^-8^<p<1×10^-6^) using data from the first and second rounds of a large schizophrenia meta-analysis by the Psychiatric Genomics Consortium. Results suggest that all methods have considerable (and similar) predictive accuracies (AUCs 0.64-0.71). However, predictive accuracy results obtained from the test set do not always reflect results obtained from the application to the schizophrenia meta-analysis. In conclusion, a variety of algorithms and annotations seem to have a similar potential to effectively enrich true risk variants in genome scale datasets, however none offer more than incremental improvement in prediction. We discuss how methods might be evolved towards the step change in the risk variant prediction required to address the impending bottleneck of the new generation of genome re-sequencing studies.

---

Complex diseases are caused by the interplay of many genetic variants and the environment, and represent a considerable health burden. Genome-wide association studies (GWAS) have had success in identifying the genetic risk factors involved in complex diseases such as inflammatory bowel disease^1^ and schizophrenia^2^. Interrogating the entire genome, exome or even selected genes through next generation sequencing technologies have also identified further risk variants^3–6^. However, more disease-associated variants, hereafter called risk variants or hits, remain to be discovered. Some risk variants are difficult to detect by current techniques due to limited sample sizes and low effect size of the variants. *In silico* methodologies that integrate evidence over multiple data sources have the potential to unearth some of these risk variants in a cost-effective manner. The novel risk variants that are identified will help illuminate the genetic risk factors involved in complex diseases, which in turn could lead to earlier or more accurate diagnoses, and the development of personalized treatment options.

Risk variants show enrichment in functional annotations, such as DNase I hypersensitive sites, transcription factor binding sites, and histone modifications (for example,^7–9^). Several groups have gone further with the results of enrichment by incorporating functional annotations as predictor variables in statistical learning frameworks to prioritize genetic variants for further study^10–12^. These statistical learning algorithms use the functional annotations to define a model that provides some measure of whether a variant is likely to increase the risk of manifesting a complex trait. However, understanding the relative merits of these approaches requires a thorough investigation into which statistical learning algorithm and/or which combination of functional annotations most effectively identifies novel risk variants.

We compared the performance of three published methods that differ in annotation set, algorithm and training data: a regularized regression called elastic net from Gagliano et al. (14 annotations)^10^, a modified random forest from Ritchie et al. (174 annotations)^12^ and a support vector machine from Kircher et al. (949 annotations, expanded from 63 unique annotations)^11^. These three papers describe algorithms capable of incorporating a large number of genetic variants labeled with multiple functional annotations, and can output a prediction score for each variant; hence, they are highly comparable.

There are many aspects to consider in the statistical learning framework (**Supplementary Fig. 1**). First, the genetic data input consists of both known risk variants and corresponding control variants (those with no evidence for risk effect). Of the variants in the genome, there is an overwhelming presence of non-risk compared to known risk variants, and this may also impact the statistical learning algorithm. Known risk variants may be identified from sources, such as the NHGRI GWAS Catalogue^13^, the ClinVar database^14^ and the Human Gene Mutation Database (HGMD)^15^. In addition, the variants can be simulated. For example, Kircher et al. used an empirical model of sequence evolution with local adjustment of mutation rates^11^. In this way, the simulated variants would contain *de novo* pathogenic mutations.

With regard to the functional annotations, some come from experimental procedures while others are predicted computationally. Examples include genomic and epigenomic annotations that can be incorporated from various online browsers and collections such as the Ensembl Variant Effect Predictor (VEP)16 and the ENCODE Project^17^.

Finally, there are numerous statistical learning algorithms from which to choose. These algorithms must be able to handle the features of the functional data: correlations among predictor variables, and a large quantity of both samples and predictor variables.

In this paper we present an investigation of variant prioritization methods that encompasses nine models: combinations of three different statistical learning algorithms and three different functional annotation sets (summarized in Table 1). The sets of nine models were created for different classifications of hits: the NHGRI GWAS Catalogue^13^ or the Human Gene Mutation Database (HGMD)^15^. HGMD contains disease variants, whereas the GWAS Catalogue contains variants associated with disease, but those variants may only be tagging the “causal” variant. However, models based on GWAS data can be tested effectively in current data (we apply those models to the schizophrenia GWAS from the Psychiatric Genomics Consortium). The HGMD models were created to attempt to create models based on next generation sequencing data, but as there are not a sufficient number of variants identified from sequencing studies to date, these models cannot yet be trained or tested with such empirical data.

**Table 1.** Comparison of the three papers.

|  | **Gagliano *et al.***<br>**(PLOS ONE 2014)** | **Ritchie *et al.***<br>**(Nat Methods 2014) “GWAVA”** | **Kircher *et al.***<br>**(Nat Genetics 2014) “CADD”** |
| --- | --- | --- | --- |
| Functional annotations | n = 14 (ENCODE, eQTLs, PhastCons, Genic context...) | n= 174 (ENCODE, GERP, Genic context...) | n = 63 (expanded to 949) (Ensembl VEP, ENCODE, PolyPhen...) |
| Risk variants (“Case”) | NHGRI GWAS Catalogue (p-value $\leq 5 \times 10^{-8}$) | HGMD - “regulatory” | Simulated mutations under neutral model - “gap” sites |
| Non-risk variants (“Control”) | union of common Illumina and Affymetrix GWAS panels | other variants in 1000 Genomes Project (for example, within 1kb of each HGMD variant) | high-frequency derived human alleles from 1000 Genomes |
| Classifier algorithm | Elastic net | Random forest | Support vector machine |
| Training protocol | 60% training<br>40% reserved for testing | 100% training | 99% training<br>1% reserved for testing |

## Results

Our primary analysis used the NHGRI GWAS Catalogue as the classifier. The test set was used to determine accuracy. These results are presented below.

### Area under the ROC curve

All the models had similar accuracy as demonstrated by the area under the curve (AUC) in the test set data (Table 2). Models using Kircher et al.’s annotations produced slightly higher AUCs compared to the other two annotation sets for the elastic net and random forest algorithms.

**Table 2.** The area under the curve (AUC) for the GWAS Catalogue comparisons, holding data and classifier constant, while varying algorithm and annotations. The AUC in the training set is in parentheses.

| **Annotations →** | **Gagliano *et al.*** | **Ritchie *et al.*** | **Kircher *et al.*** |
| --- | --- | --- | --- |
| Elastic Net | 0.67 (0.67) | 0.65 (0.67) | 0.71 (0.74) |
| Random Forest (altered minimum node size) | 0.67 (0.69) | 0.68 (0.72) | 0.70 (0.79) |
| Support Vector Machine (with prior feature selection) | 0.66 (0.66) | 0.64 (0.66) | 0.64 (0.68) |

The AUC results for the training set were also computed to investigate whether the models were over-fit; that is to say, whether the training set AUC is much higher than the test set AUC. We found that for the 174 and 949 annotation sets, the random forest models with node size equal to one were prone to over-fitting. For instance, for the random forest model based on the 174 annotations, the test set and training set AUCs were 0.687 and 0.998, respectively. The over-fitting in the random forest models was solved when the minimum node size was set to 10% of the total data size. These results highlight the importance of ensuring that appropriate parameters are chosen for the algorithms. Due to the over-fitting, only the random forest models with the minimum node size equal to 10% of the data will be discussed in the rest of the results.

### Density and distribution of prediction scores

Violin plots were constructed by plotting the prediction scores for hits (risk variants) and non-hits in order to visualize how well the two classes were separated (Fig. 1 and **Supplementary Table 1**). The distributions for the hits and non-hits overlapped for all the models. Models with the highest AUCs did not always show the better separation between the hits and non-hits according to the violin plots. These results suggest that the AUC may be driven by the correct identification of the larger class (non-hits), regardless of performance for identifying the hits. The two models with the best AUCs (Kircher et al. annotations with elastic net (0.71) and with random forest (0.70)) have well separated means and relatively normal distributions. In one of the two models with the lowest AUC (Ritchie et al. annotations with support vector machine (0.64)), the median prediction score between hits and non-hits is most similar and the distribution is very skewed. Interestingly one of the mid-range performance models, the Gagliano et al. annotations for the support vector machine (0.66) shows evidence of a multimodal distribution where one mode is more common in the hits and another in the non-hits. However, this effect may simply be due to the comparatively small number of annotations, which lead to a smaller number of possible scores. Generally, the models created using the Kircher et al. annotations showed the largest spread of the prediction scores for both the hits and non-hits.

**Figure 1.**
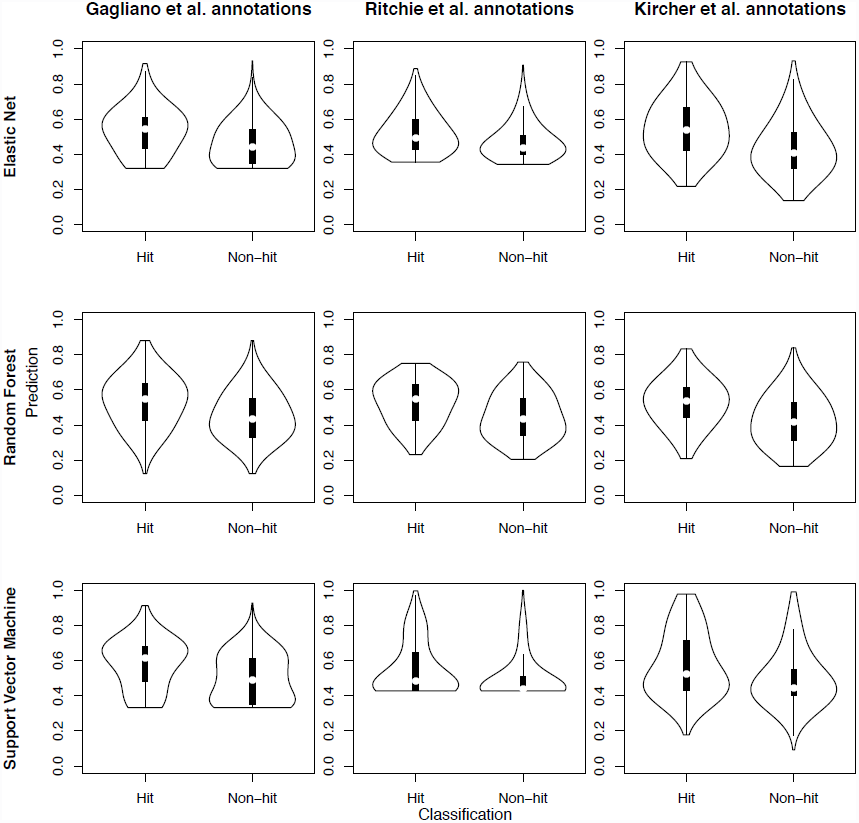
Violin plots showing class separation by prediction scores for the various comparisons using the GWAS Catalogue as the classifier. The non-scaled elastic net models are plotted. The adjusted minimum node size random forest models are plotted.

### Feature selection within elastic net and random forest

More does not necessarily equal better as not all the annotations may be relevant to predicting risk variants. Generally, not all of the functional annotations in the annotation sets (N=14, N=174 and N=949) were used to create the various models. For instance, elastic net assigned non-zero Beta coefficients to 9 out of 14 annotations, 12 out of 174, and 16 out of 949. Random forest assigned non-zero Gini importance values to all of the 14, 131 out of 174, and 239 out of 949. All of these models had similar performance in the test sets (AUCs ranging from 0.68 to 0.70 for the random forest models and 0.65 to 0.71 for the elastic net models). Some of the features in the training set were invariable, or in other words none of the variants had the annotation. Of the 14 annotations in Gagliano et al., none were invariable; of the 174 annotations in Ritchie et al., three were invariable; and of the 949 annotations in Kircher et al., 517 were invariable. The results suggest that elastic net has a more stringent feature selection implementation than random forest. The support vector machine models always assigned non-zero feature weights, as support vector machine does not intrinsically perform feature selection, as does elastic net and random forest. Thus, we inputted only those annotations with a non-zero Beta coefficient from the elastic net models into the support vector machine models (see **Methods**).

### Importance of the functional annotations

Different combinations of annotations can be used to obtain models with similar predictive accuracy. However, it is difficult to interpret the importance of the annotations for numerous reasons, some of which are discussed below.

All three annotation sets contained a mixture of binary variables and continuous variables. For Kircher et al.’s annotations, background selection (the annotation with the widest continuous scale that ranged from 0 to 1000) came up as most important for predicting the class label in the random forest model. This bias for random forest preferentially selecting annotations measured on a continuous scale has been previously described^18^. When making a decision at a node, continuous annotations can be used multiple times at varying cut-offs to split the data. In this way, functional annotations measured on a continuous scale are incorporated more often into the forest compared to non-continuous annotations, and thus obtain higher variable importance measures^18, 19^.

It is also difficult to interpret the variable importance measures derived from elastic net because this algorithm is not scale invariant and the annotations were not standardized. Random forest, however, does not require an additional step to scale the annotations prior to incorporating them into the model.

Using Gagliano et al.’s annotations with elastic net, we compared the models created with scaled (all annotations have a standard deviation of 1 and a mean of 0) versus non-scaled annotations. Although the AUCs for both models were nearly identical, the assigned Beta coefficients differed (**Supplementary Fig. 2a**). The assigned Gini importance measures for the random forest models are shown in **Supplementary Figure 2b**, and the weights assigned to the variables in the support vector machine model using only those features that had non-zero Beta coefficients in elastic net are in **Supplementary Figure 2c**. See **Supplementary Files 1-3** for the feature importance measures from all the models based on the GWAS Catalogue as the classifier.

### Performance for complex disease variants: Application to Schizophrenia GWAS

Various quantile-quantile plots were constructed in order to compare which models showed greater separation of the schizophrenia GWAS p-values for high scoring and low scoring functional variants. Plots were constructed where annotations were held constant but the algorithm differed (Fig. 2). For instance, for the 14 annotations from Gagliano et al. we plotted the models from the three algorithms in one plot. Furthermore, models from the same algorithm but varying by annotation set were compared (**Supplementary Fig. 3**). With regard to the functional annotation set, the separation of the novel associated variants from the non-associated in the sub-genome-wide-significant variants was best exhibited when using either the Kircher et al. or Ritchie et al. annotation sets. Regardless of annotation set, the elastic net models consistently showed good separation. The results for the application to the schizophrenia GWAS did not always reflect the AUCs. For instance, a poor performing model in terms of AUC based on the test set, elastic net with the Ritchie et al. annotations, performed well in the GWAS application. All in all, the accuracy of the resulting models should be assessed by various means, including (but not limited to) theoretical models such as the ROC curve, as well as empirical approaches such as applying the model using data from one study and evaluating its performance on independent data with gold standard answers.

**Figure 2.**
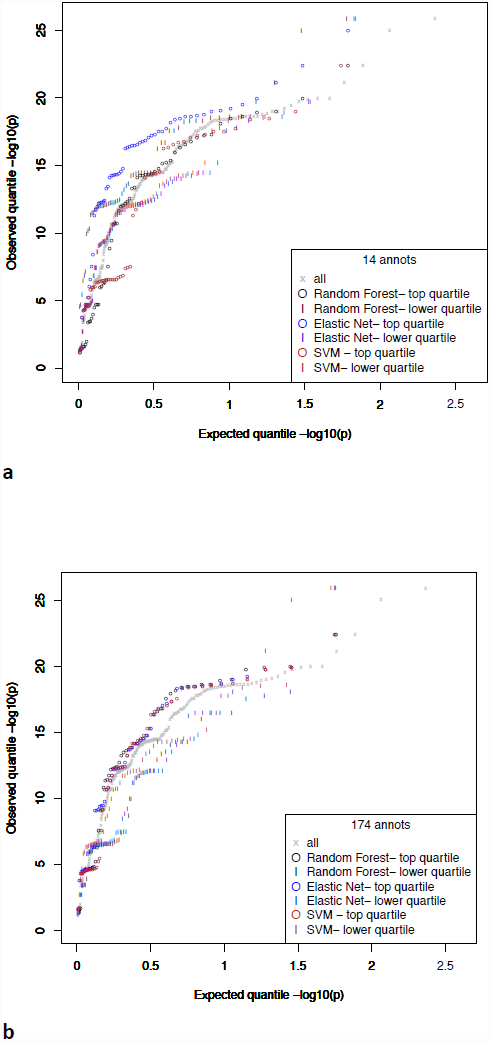
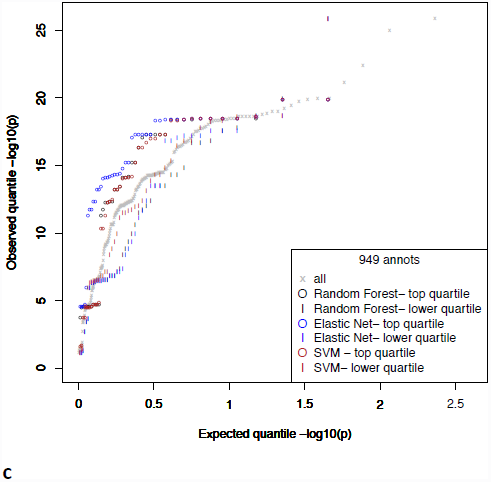
Quantile-quantile plots of PGC1 sub-genome-wide-significant variants (5×10-8<p<1×10-6) stratified by prediction score for the various models based on the GWAS Catalogue classifier, and plotted by PGC2 p-values. Models grouped by annotation set: 14 [**a**], 174 [**b**], and 949 annotations [**c**]. The lower quartile genetic variants are those with a prediction score in the first quartile, and the top quartile variants are those with prediction values in the fourth quartile.

### HGMD Analysis

The models for the analysis using all of the HGMD variants using either the Ritchie et al. or Kircher et al. annotations had high predictive accuracy (Table 3). The AUCs for the non-exonic HGMD analysis were more comparable to the ones obtained for the primary analysis using the GWAS Catalogue as the classifier (Table 4), but again the annotations from Ritchie et al. and Kircher et al. performed better.

**Table 3.** The area under the curve (AUC) for the HGMD comparisons, holding data and classifier constant, while varying algorithm and annotations. The AUC in the training set is in parentheses.

| **Annotations →** | **Gagliano *et al.*** | **Ritchie *et al.*** | **Kircher *et al.*** |
| --- | --- | --- | --- |
| Elastic Net | 0.66 (0.65) | 0.87 (0.88) | 0.88 (0.88) |
| Random Forest (altered minimum node size) | 0.65 (0.66) | 0.91 (0.91) | 0.87 (0.89) |
| Support Vector Machine (with prior feature selection) | 0.61 (0.66) | 0.85 (0.86) | 0.85 (0.87) |

**Table 4.** The area under the curve (AUC) for the non-exonic HGMD comparisons, holding data and classifier constant, while varying algorithm and annotations. The AUC in the training set is in parentheses.

| **Annotations →** | **Gagliano *et al.*** | **Ritchie *et al.*** | **Kircher *et al.*** |
| --- | --- | --- | --- |
| Elastic Net | 0.65 (0.66) | 0.77 (0.78) | 0.79 (0.80) |
| Random Forest (altered minimum node size) | 0.65 (0.65) | 0.80 (0.86) | 0.78 (0.85) |
| Support Vector | 0.61 (0.68) | 0.68 (0.78) | 0.76 (0.82) |
| Machine (with prior feature selection) |  |  |  |

Similar to the analysis using the GWAS Catalogue as the classifier, for the HGMD analysis models the features that came up as most important tended to vary depending on the algorithm. Genic annotations featured highly (see **Supplementary Files 4-6)**. For the Gagliano et al. annotations, the top annotation (or the second most important in the case of support vector machine) was nonsynonymous SNPs. For the Kircher et al. annotations the top annotations for the random forest and support vector machine models were related to the coding sequence or nonsynonymous SNPs. For instance, the top two annotations for support vector machine were nonsynonymous SNPs, and the interaction between nonsynonymous SNPs and minimum distance to the transcribed sequence end (TSE). The top for random forest were amino acid position from coding start (“protpos”), the interaction between nonsynonymous SNPs and base position from coding start, etc. Finally, the top annotation for elastic net was CpG. For the Ritchie et al. annotations, the top two were coding sequence and exon for both the random forest and support vector machine models. For elastic net, the top two were donor and coding sequence. The importance of genic features is likely linked to bias in the data, which will be examined further in the discussion.

The HGMD analysis in which only non-exonic HGMD and control variants were considered seemed to overcome this bias towards genes or positions relative to genes. Features related to genes were not consistently coming up as most important for all the models. In fact, for all algorithms, the top annotation for the Gagliano et al. annotation set was DNase I hypersensitive sites (see **Supplementary Files 7-9**).

## Discussion

We found that the three algorithms assessed here, elastic net, random forest and linear support vector machine show comparable accuracy in GWAS data. Furthermore, our results show that various combinations of annotations can create models with similar predictive ability when it comes to identifying risk variants from non-risk variants. This observation makes it difficult to differentiate the predictive power of the functional annotation sets used by each study, at least in the case of GWAS risk variants.

To apply the methods in next generation sequencing data we would ideally use risk variants identified from such studies. Unfortunately, there are not a sufficient number available. We used the HGMD to attempt to extrapolate our findings. However, we believe the high accuracies achieved for the all HGMD models (ie. not just the models looking at non-exonic variants) are driven by the inherent bias of the HGMD data, in that it is largely focused on genes. For the models using only non-exonic HGMD and control variants, the AUCs were considerably lower, with the Kircher et al. and Ritchie et al. annotation sets clearly out-performing the annotations used by Gagliano et al. Yet, this subset is a highly derived and filtered set of variants, emphasizing the need for empirical data. There are even some issues with the GWAS hits used for our primary comparisons where we identified risk variants as those variants that appear in the NHGRI GWAS Catalogue^13^ with a p-value of ≤ 5×10^-8^: GWAS variants may only be tagging the actual “causal” variants. The simulation employed by Kircher et al., in which the functional annotations were used to differentiate between millions of high frequency human-derived alleles from the same number of simulated alleles,^11^ showed considerable accuracy; further adaptions to this strategy may prove useful. Ideally, we require empirical data from actual next generation sequencing studies, but to date there are not a sufficient number of novel complex trait genetic risk variants identified through this approach to train or test the models.

The support vector machine models did not perform as well as their corresponding elastic net or random forest models as exhibited by lower AUCs for the support vector machine models for the GWAS Catalogue and all HGMD analyses. This poor performance may be attributed to the fact that we implemented the most basic kernel type for the support vector machine, a linear kernel. This kernel was chosen to reduce computational time, and also in an effort to be consistent with the type of kernel that was utilized by Kircher et al. A linear kernel may not be best to separate the data. Furthermore, as support vector machine does not intrinsically perform feature selection, we selected those features with a non-zero Beta coefficient from the corresponding analysis using the elastic net algorithm. Use of another method of feature selection may have yielded different results. Our results do not necessarily suggest that the elastic net and random forest algorithms out-perform the support vector machine algorithm, since altering either the kernel type or the functional annotations in the support vector machine models may produce results comparable to the other two algorithms.

There are limitations to this comparison. For example, other statistical learning algorithms and other annotation sets could be explored. Annotation sets could be phenotype specific, as there is evidence that the level of enrichment of functional information can differ depending on the subset of risk variants selected^20^. For instance, enrichment of disease-specific variants in the GWAS Catalogue can differ in certain cell types, for example for DNase I hypersensitive sites^8^.

Identifying which algorithm and/or annotations identify risk variants with the highest accuracy will help researchers develop a better understanding of the genetic factors involved in complex disease in a cost-effective manner making use of a rich set of publically available functional data. This work helps illuminate the genetic factors involved in disease by making use of existing functional data *in silico*. Increasing knowledge on the etiology of complex disease will allow for earlier or better diagnoses, and the development of personalized treatment and novel therapies.

## Methods

We explored the utility of each of the three algorithms with each of the three functional annotation sets in order to attribute performance differences to the algorithm and/or annotations. A total of nine models were created.

In the primary analysis, the set of risk variants used for training all the models were based on whether or not a genetic variant is a hit or a non-hit from a genome-wide association study (GWAS). Hits were defined as those variants present in the NHGRI GWAS Catalogue (www.genome.gov/gwastudies, downloaded on August 7, 2014)^13^ with a p-value of equal to or less than 5×10^-8^. There were 3,618 unique genetic variants that met these criteria. (Note that at the time of download the novel hits from the second phase of the schizophrenia GWAS from the Psychiatric Genomics Consortium (PGC2)2 had not yet been included.) A subset of non-hits was selected from common GWAS arrays (Affymetrix Genome-Wide Human SNP Array 6.0, the Illumina Human1M-Duo Genotyping BeadChip, and the Illumina HumanOmni1-Quad BeadChip). Those non-hits in high linkage disequilibrium (r2>0.8) with hits were removed from the analyses.

### Functional annotation sets

The data was then annotated using three distinct protocols outlined in each of the three respective papers. The variants were marked with the Gagliano et al. annotations available on the website (http://www.camh.ca/en/research/research_areas/genetics_and_epigenetics/Pages/Statistical-Genetics.aspx). Fourteen functional annotations were used by Gagliano et al., two of which were on a continuous scale (two conservation measures, PhyloP and PhastCons), and the remaining were binary, signifying the presence or absence. The binary annotations included those related to genomic context such as the presence in a gene, a splice site or a transcription start site, as well as those from the ENCODE Project^17^ such as three types of histone modifications and DNase I hypersensitivity. For the ENCODE data, functional annotations present in multiple cell lines were grouped together, and genetic variants were annotated accordingly in a binary, present or absent, fashion.

To annotate the variants using Ritchie et al.’s annotations, the data were entered into the online GWAVA webserver (https://www.sanger.ac.uk/resources/software/gwava/). Ritchie et al. investigated 174 functional annotations, some binary and others continuous. They also used ENCODE Project tracks including those investigated in Gagliano et al. but not necessarily coded as presence or absence. For instance, for transcription factor binding sites, the number of cell types in which the site was present was used as the annotation. Additionally, human variation (for instance mean heterozygosity) and genic and sequence contexts (for instance gene region annotations provided by Ensembl, and GC content, respectively) were included as well.

To obtain Kircher et al.’s annotations, the data were entered into the online CADD webserver (http://cadd.gs.washington.edu). However, Kircher et al. also imputed missing values, expanded categorical variables, added indicator variables, and included interaction terms. Martin Kircher provided scripts to run on the webserver output to prepare our dataset in accordance with the complete protocol. Kircher et al. looked at 63 unique functional annotations, which totaled to 949 once the categorical variables were expanded, and the indicator variables and interaction terms were included. A mixture of continuous, categorical, and binary functional annotations was included. Similar annotations to those used by Gagliano et al. and/or Ritchie et al. were included, such as ENCODE Project annotations and genic context. Additionally, data from online variant prediction programs (e.g. Sift^21^ and PolyPhen^22^) were incorporated.

### Statistical learning algorithms

The variants were randomly divided; 60% was used for training the models, and the remaining 40% was reserved for testing. Elastic net is a regularized logistic regression, which by the use of penalty parameters prevents the coefficients from getting too large. The elastic net models were constructed using the glmnet package in R^23^. A weighting procedure was included to up-weight hits, as described in Knight et al.^24^; in brief, the weighting has the effect of equalizing the number of hits and non-hits in the training set. Optimal parameters of lambda and alpha were selected for each elastic net model. Lambda is an overall penalty parameter. Alpha controls the proportion of weight assigned to both the sum of the absolute value of the coefficients and the sum of the squared value of the coefficients, which affects the degree of their sparsity. A range of combinations of lambda and alpha were investigated. The lambda and corresponding alpha that give a model a deviance one standard deviation above the model with the lowest deviance was selected.

Random forest is a collection of decision trees. The random forest models were implemented in Python using the scikit-learn package^25^. Two sets of random forest models were created. For the first set, we replicated Ritchie et al.’s random forest implementation by using scripts (e.g. gwava.py) provided on their online GWAVA FTP site (ftp://ftp.sanger.ac.uk/pub/resources/software/gwava/). For instance, bootstrap sampling was employed to form decision trees from bootstrap subset samples. To address the class imbalance in the datasets, non-hits were down-weighted through the balance_classes function created by Ritchie et al. and included in their random forest implementation. The balance_classes function selects a subset of non-hits that is equal to the number of hits in order to grow a tree. Furthermore, the subset of annotations used to determine the node split was set to the square root of the total number of annotations. This setting is the default setting for classification problems to determine the best split at each node of the decision tree^26^. Additionally, as done by Ritchie et al., we used 100 decision trees since we determined that the prediction scores and variable importance measures did not significantly differ past 100 trees.

Ritchie et al. used a minimum node size (min_samples_split) of 1. The minimum node size is the minimum number of samples required to split an internal node. We created another set of random forest models in which we adjusted the minimum node size. This parameter is dataset specific, and a recommended setting is 10% of the total dataset^26^. Consider n to be the number of hits in the training dataset. For the second set of random forest models, we set the minimum node size to approximately 10% of 2n.

Support vector machine creates a hyperplane within a decision boundary space defined by support vectors to separate the classes in multidimensional space. The support vector machine models were implemented in Python through the scikit-learn package^25^. Kircher et al. did not use a weighting procedure as their training set was already balanced. To compare protocols in an unbiased manner, we used a subset of the training set in which we chose all hits, and randomly selected an equal amount of non-hits. We performed a grid search using the tune function in order to determine the optimal cost parameter for a linear kernel. The cost parameter is a penalty (see chapter 9 in James et al.^27^ for details). Feature selection is critical to improving model performance and is intrinsically incorporated by the elastic net and random forest algorithms^28^. Feature selection must be implemented before using support vector machine, as there is no feature selection protocol built in. Kircher et al. utilized univariate logistic regression among other methods to select features that best predict genetic risk variants. In this paper our support vector machine models included those annotations that had a nonzero Beta coefficient from the corresponding elastic net models. We chose the annotations found to be important from elastic net, since this algorithm implements a more stringent feature selection protocol compared to random forest (see **Results**).

### Assessment of model performance

We assessed model performance in the test set data by calculating the area under the receiver operating characteristic (ROC) curve. As another measure of model performance, we also examined the distribution of prediction scores assigned to the test set data with the aid of violin plots. Furthermore, we investigated importance of the functional annotations through the Beta coefficient for elastic net. Similar to the output from a simple logistic regression, the larger coefficients are interpreted as more important to predicting genetic risk variants. For random forest we used Gini importance, which was also used in Ritchie et al. Gini importance is a scaled measure of Gini impurity averaged over all trees; it represents the improved capacity for correctly predicting variants that can be directly attributed to the annotation^29^. For support vector machine, feature weights can be obtained related to the construction of the hyperplane when a linear kernel is used^30^.

### Performance for complex disease variants: Application to Schizophrenia GWAS

We tested the performance of the nine models based on the GWAS classifier in a schizophrenia GWAS context. We selected all sub-genome-wide-significant variants (5×10^-8^<p<1×10^-6^) from the first round of the GWAS by the Psychiatric Genomics Consortium (PGC1)31. For each of the nine models we obtained prediction scores for these variants and selected the variants from the first and fourth prediction score quartiles. For these variants we extracted the p-values from the larger second round of the GWAS (PGC2)^2^ and plotted these in quantile-quantile plots. We were able to determine for all models whether variants assigned higher scores were enriched in the variants with more significant p-values compared to variants with less significant p-values.

### HGMD analysis

In an attempt to apply the algorithms and annotation set combinations to whole genome sequencing data rather than just GWAS, a different classifier was used to identify hits and non-hits, the Human Gene Mutation Database (HGMD). We conducted two analyses with subsets of the public release of HGMD provided to Ensembl in the fourth quarter of 2013 (provided by Graham Ritchie). In the first, we took all the variants (single nucleotide polymorphisms) in HGMD (N= 3,391) and chose controls that fell within a kilobase of either side from the HGMD variant. Secondly, models based on the subset of non-exonic HGMD variants (N= 689) and non-exonic control variants were assessed. This second set of models was created in an effort to overcome the ascertainment bias inherent in HGMD related to genes.

The nine models created by combinations of annotation sets and algorithms were assessed for the two sets of HGMD models described above. Additionally, the data was randomly split into 60% for training and 40% for testing. The same procedures for elastic net, random forest and support vector machine used in the GWAS Catalogue analysis were also conducted for the HGMD analyses.

### Comparison of scores from the three papers: Application to Schizophrenia GWAS

In the effort for a more general comparison of the published methods as is, rather than looking specifically at the algorithm and annotations as done above, we additionally conducted the schizophrenia GWAS application using scores for the variants obtained directly from the published papers. This analysis is further described in **Supplementary Text** and the results are depicted in **Supplementary Figure 4**.

## Acknowledgements

Computations were performed on either the CAMH Specialized Computing Cluster (SCC) or the General Purpose Cluster (GPC) supercomputer at the SciNet HPC Consortium^32^. SciNet is funded by: the Canada Foundation for Innovation under the auspices of Compute Canada; the Government of Ontario; Ontario Research Fund - Research Excellence; and the University of Toronto. The CAMH SCC is funded by: The Canada Foundation for Innovation, Research Hospital Fund. We thank David Rotenberg, and Jon Pipitone, SCC Administrators, for providing technical support. We thank Kathleen Askland, Nickhil Bhagwat and Malgorzata Maciukiewicz for support with the statistical learning. We thank Martin Kircher for support with CADD, and for providing scripts to expand the 63 annotations to 949. We also thank Graham Ritchie for support with GWAVA.

SAG is funded by the Peterborough K.M. Hunter Graduate Studentship, and is also funded through the Armstrong Family via the CAMH Foundation. SAG is a fellow of CIHR STAGE (Strategic Training for Advanced Genetic Epidemiology) – CIHR Training Grant in Genetic Epidemiology and Statistical Genetics. This research was supported by funding from Training grant GET-101831. RR was funded through the Institute of Medical Science Summer Undergraduate Research Program. The authors acknowledge financial support from the Department of Health via the National Institute for Health Research (NIHR) comprehensive Biomedical Research Centre award to Guy’s & St. Thomas’ National Health Service (NHS) Foundation Trust in partnership with King’s College London and King’s College Hospital NHS Foundation trust. JK was previously funded by this source. JK is currently funded by charitable donors at the CAMH Foundation: http://www.supportcamh.ca. JK holds the Joanne Murphy Professor in Behavioural Science. MEW is funded by the Higher Education Funding Council for England (HEFCE): http://www.hefce.ac.uk/. MRB is funded by the National Institutes for Health Research (NIHR), and this work forms part of the portfolio of translational research of the NIHR Biomedical Research Unit at Barts.

